# Genetic selection for residual feed intake impacts the functional redundancy of the gut microbiota in pigs during the growing-finishing period

**DOI:** 10.1101/2024.06.05.597549

**Authors:** Olivier Zemb, Caroline Achard, Jordi Estellé, Laurent Cauquil, Catherine Denis, Yvon Billon, Sylvie Combes, Claire Rogel-Gaillard, Hélène Gilbert

## Abstract

**Background:** Improving feed efficiency is an unvarying goal in livestock production. The role of gut microbial communities in feed efficiency was investigated in two divergent pig lines selected for their residual feed intake (RFI), which is the difference between the observed feed consumption of an animal and that predicted for its maintenance and production requirements. Hence, RFI is a measure of net feed efficiency.

**Results:** By sequencing the 16S rRNA genes from fecal samples, we found that the microbiota of low RFI pigs (LRFI; n=12) was significantly depleted in four operational taxonomic units (OTUs) belonging to *Prevotella*, *Feacalibacterium* and *Lactobacillus* genera by comparison to high RFI pigs (HRFI; n=11). Inferring KEGG orthologs from the 16S rRNA gene sequencing data showed that LRFI pigs have 20% more functional redundancy than HRFI pigs. The inferred orthologs held more discriminative power than the OTU approach, which might be inherently flawed if functionally equivalent OTUs compete for the same niche.

**Conclusions:** Gut microbiota of LRFI pigs had different microbial operational taxonomic units and a higher functional redundancy, which were more shared suggesting a role of the gut microbiota in the global feed efficiency according to Tax4Fun but this inference based on 16S data has to be verified in future work.

## Background

Improving feed efficiency to reduce feed cost is an incessant challenge for livestock production, including the pig breeding sector. Feed efficiency can be estimated by individual residual feed intake (RFI). The RFI is the difference between the observed feed consumption of an animal and the one predicted for maintenance and production requirements. It is interpreted as a net feed efficiency indicator, as compared to feed conversion ratio (FCR) which is highly correlated to production and maintenance. Comparing pig lines selected for low RFI (LRFI, more efficient) and high RFI (HRFI, less efficient) have already shown that low RFI is linked to decreased daily feed intake (DFI) [1], basal metabolic rate [2, 3], body fat [1], lipid metabolism in adipose tissue [4, 5], glycogen content in muscle [6], and changes in catabolic and oxidative pathways in the muscle, the liver and the adipose tissue [7-9]. It has already been reported that selection for improving feed efficiency during growth with standard highly digestible diets or with diets enriched in dietary fibers had no impact on digestibility in either low or high RFI lines (Barea *et al.* [2] and Montagne *et al.* [10]).

The microbial communities residing in the digestive tract are known to play a main role in the digestion processes, absorption of nutrients and metabolism [11], but very little is known on their role in the variation of feed efficiency in pigs. Our aim was thus to characterize the gut microbiota of high (HRFI) and low RFI (LRFI) pig lines by sequencing the 16S rRNA gene of fecal DNA samples, and to investigate i) whether specific microbes were characteristic of LRFI pigs, and ii) whether specific microbial functions were different between the two pigs lines, even if they are shared by multiple microbial species.

## Methods

### RFI lines

A divergent selection program on RFI in growing pigs started in 2000 [1]. Briefly, purebred French Large White pigs were divergently selected on RFI to constitute two pig lines referred to as the LRFI and HRFI lines. The animals were housed in INRA experimental farms at Rouillé and Le Magneraud, France. The piglets were weaned at 28 days old. They further gathered at the Rouillé INRA and penned by groups of 24 per line. At 10 weeks of age, male candidates to selection were moved to the growing-finishing unit and penned by groups of 12 per line without mixing of the post-weaning groups. In the present study, one pen with 12 male candidates of the 7th generation of selection was tested per line. Phenotypic performances were measured on animals between 35 kg body weight (BW) and 95 kg BW. One pig of the HRFI line died during the test and was excluded from the analyses. Pigs were weighted individually every week to define the days when they reached 35 and 95 kg BW, and average daily gain (ADG, in grams per day) was computed for this test period. Daily feed intake (DFI, in grams per day) was individually recorded with automatic self-feeders ACEMA 64 (ACEMO, Pontivy, France). At 95 kg BW, ultrasonic backfat thickness (UBT, in millimeters) was measured as the average of six measures (right and left shoulder, right and left midback, right and left loin) (ALOKA SSD-500 echograph; Aloka, Cergy Pontoise, France). In order to account for growth requirement, an RFI selection index in grams per day was computed as a linear combination of DFI, ADG and UBT traits as follows [1]: RFI= DFI - (1.06 x ADG) - (37 x UBT) + 100.

With this computation, pigs with RFI lower than 100 were more efficient than on average and pigs with RFI higher than 100 were less efficient than on average. In generation G7, the LRFI and HRFI lines differed by 136.8 g/d of RFI, i.e. more than 3.1 genetic standard deviations of the trait [12]. Pigs were offered free access to water. A pelleted commercial diet based on cereals and soybean meal containing 10 MJ NE/kg and 160 g CP/kg, with a minimum of 0.80 g digestible Lys/MJ NE formulated for growing pigs was offered *ad libitum* to the pigs (Table S1). No disease was observed during the test period and the pigs were not fed antibiotics.

### Growth and feed ingestion measurements and stool fecal sampling

In addition to routine measurements for selection on RFI as described above, the 23 pigs were weighed at 11 (74.3 ± 1.1 days), 17 (116.3 ± 1.1 days) and 20 weeks of age (137.3 ± 1.1 days). Fecal samples were collected during these three intermediate weighing and immediately placed in dry ice for storage at -80°C before DNA extraction. Differences in performances between LRFI and HRFI pigs were assessed using the Student test, with a model including the effect of the line only.

### Analysis of the microbiota composition

Fecal samples (n=69) were kept at -80°C until DNA extraction. The DNA was extracted from 200 mg of porcine stool using a bead-beating approach as described previously [13], with a protocol that has been recently validated by the International Human Microbiome Standards (IHMS) project [14]. The V3-V4 region of the gene coding for the 16S rRNA gene was amplified, tagged and pooled in one single sequencing run of a Roche GS FLX Titanium instrument [15]. The sequences were submitted to NCBI’s SRA archive [16] with accession number <u>SRP090793</u>.

The resulting 1.3 M sequences were filtered for quality (length>150bp, homopolymers<10, 100% unambiguitous bases) and reassigned to each sample using barcodes, which were also trimmed. The chimeras were removed using the Uchime algorithm [17], clustered into operational taxonomic units (OTU) using ESPRIT-TREE [18] and counted in each sample to create a table of relative abundance of each OTU across the samples, yielding between 1144 and 2726 high-quality sequences per sample. The shannon index (vegan R package) was analyzed with an age x line linear model.

To assess differences in OTUs relative abundance between groups, a zero-inflated model implemented in the fitFeatureModel function of the metagenomeSeq R package v1.16.0 [19] was used. Correction for multiple testing was performed using the Benjamini-Hochberg procedure.The global dissimilarity in microbial composition was tested using the ADONIS test on the Bray-Curtis distances (vegan package, R software). These distances were represented by non Metric Dimensional Scaling. The OTU table of abundance was normalized by the number of reads per sample and used to discriminate the RFI lines using principal component analysis and discriminant analysis (DAPC) [20]. Briefly, in DAPC the samples are described in orthogonal axes using a principal component analysis and a discriminant analysis is applied on the first few components. The contribution of each OTU is then computed by multiplying the projection coefficients. The minimal number of OTUs to discriminate the groups was determined to obtain 95% of the area under the receiver operating characteristic (ROC) curve. The significance of the relative abundance of selected OTUs was tested using a Kruskal-Wallis test. The ability to separate LRFI and HRFI pigs based on their microbiota was computed by calculating the area under the ROC curve (pROC package, R software). The non-linear relationships between the selected OTUs were characterized by the maximal information coefficient (MIC) index [21]. To account for the background noise of MIC indexes, the MIC was computed in 1000 random permutations of the relative abundances within the variables and MIC values from the real dataset had to be higher than 99% of the MIC values from the randomized dataset (i.e. 0.38233) in order to be considered significant.

### Prediction of microbiota KEGG ortholog functions

The KEGG orthologs (KO) for each fecal DNA sample were reconstructed from the genus-resolution table obtained by SilvaNGS at 98% identity [22] with the Silva 123.1 database [23] using Tax4Fun with the normalization by the number of copies of the 16S rRNA genes [24]. Briefly, this software maps the table of OTU created by SilvaNGS to the 4111 microbes underlying the KEGG database to create a functional KO profile of each sample. The predicted KOs were then organized in metabolic pathways using ipath2 [25] and KOs that were unambiguously mapped on metabolic pathways were compared between the lines using a Wilcoxon test adjusted with the Benjamini Hochberg procedure [26]. The contribution of each OTU to the taxonomic profile of the samples was evaluated through an *in-silico* enrichment of the considered OTU and subsequent analysis of the taxonomic profile. When using DAPC on the KO data, the 9 selected KOs were tested using a Kruskal-Wallis test.

## Results and discussion

### The LRFI and HRFI pigs differed in performances

As expected, the LRFI pigs tended to have lower RFI than HRFI pigs (91.9 ± 24.0 g/d vs 107.2 ± 11.6 g/d, respectively, p=0.07) (Fig 1). The LRFI pigs had also lower DFI (1941 ± 229 vs 2242 ± 170 g, p<0.001), lower ADG (839 ± 50 vs 923 ± 70 g/d respectively, p<0.004) and a thinner backfat layer (13.3 ± 1.4 vs 14.9 ± 1.5 mm, p<0.002). Average BW was 29.0 ± 3.9 kg at 11 weeks of age and did not differ between lines (p=0.87). The average BW was significantly different from 17 weeks onwards (see Table S2). Considering the limited number of animals, these differences were as expected based on our previous studies (Gilbert et al, 2017).

**Figure 1:**
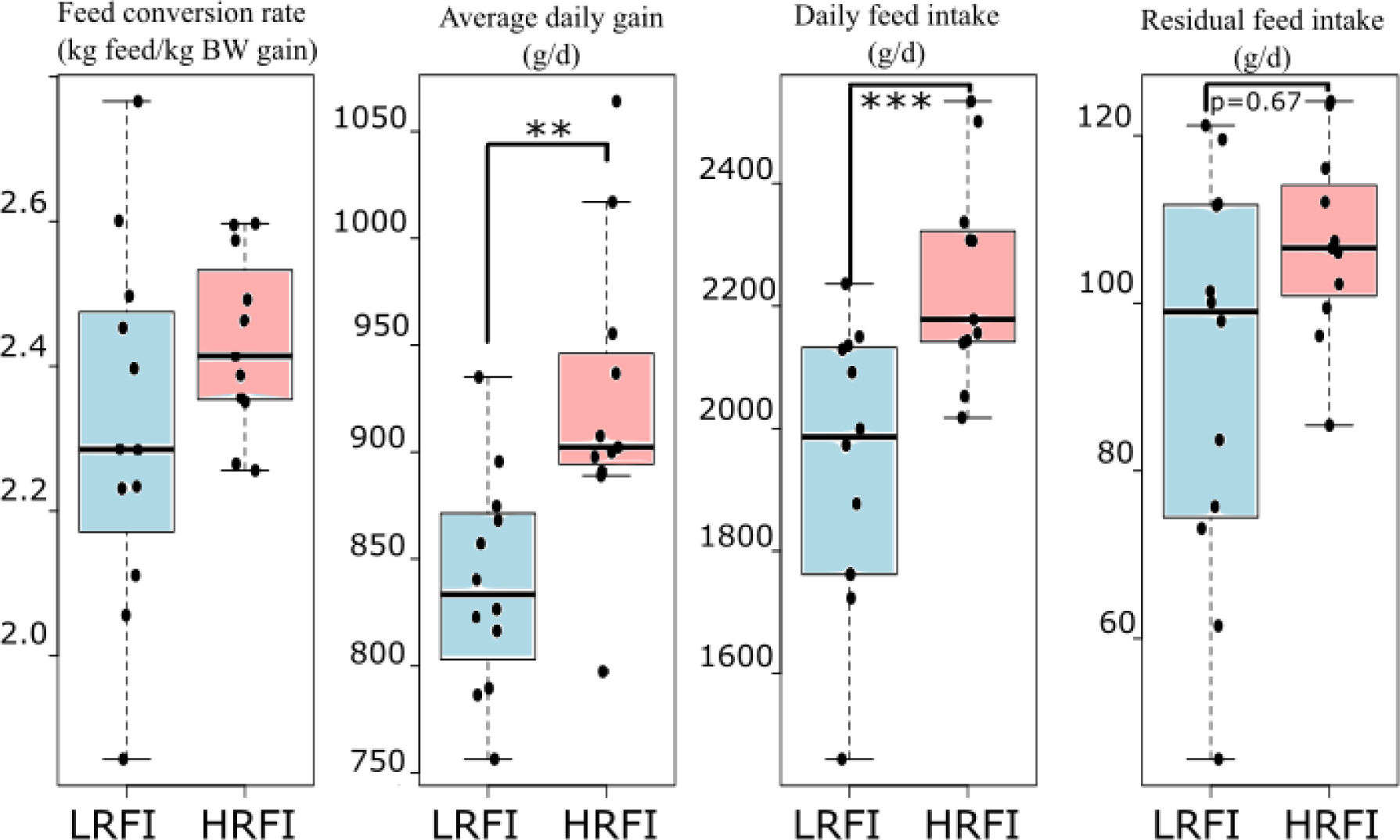
Growth parameters and residual feed intake of low RFI (in blue) and high RFI (in red) of pigs between 35 and 95 kg. ***, p=0.001, **, p=0.004

### LRFI and HRFI lines have different OTU patterns

The microbial communities contained 26573 OTUs across the 69 samples, 9977 of which were not singletons. Applying the 0.005% filter [27] to the OTU table reduced the diversity to 1272 OTUs. As expected, the major phyla was *Bacteroides* (59%) and the most abundant genera was *Prevotella* (44%) [28], while 34% of the sequences remained unclassified at the genus level. The Shannon index of the microbial communities did not vary between the lines even though it increases as pigs are aging (3.2 at 11 weeks, 3.7 at 17 weeks and 4.0 at 20 weeks, p<10^-6^).

Using the Bray-Curtis distance, both the interaction effect between line and the time of sampling was significant (p<0.001): Microbial communities from LRFI and HRFI were significantly different at 11 and 17 weeks but not at 20 weeks (p_ADONIS_<0.06, 0.002 and 0.138, respectively, Figure 2). The Metagenomeseq differential abundance analysis and adjusting for the false discovery rates only obtained two OTUs that differed between the lines despite the time effect: OTU25577 belonging to *Lactobacillus* were 10 fold less abundant in LRFI pigs than in their HRFI counterparts (0.03 % vs 0.4 % respectively). The closest cultivable representative to OTU25577 is the mucus-binding *Lactobacillus reuteri* ATCC53608 (ID 97%), which has been isolated from the pig intestinal tract [29]. The *Lactobacillus*-related OTU26000 varied between 0.4 and 1.8 % in LRFI vs HRFI pigs and shared 99% identity with *Lactobacillus johnsonii* Strain N6.2, a homofermentative lactic acid intestinal bacterium isolated from rat’s stool [30].

**Figure 2:**
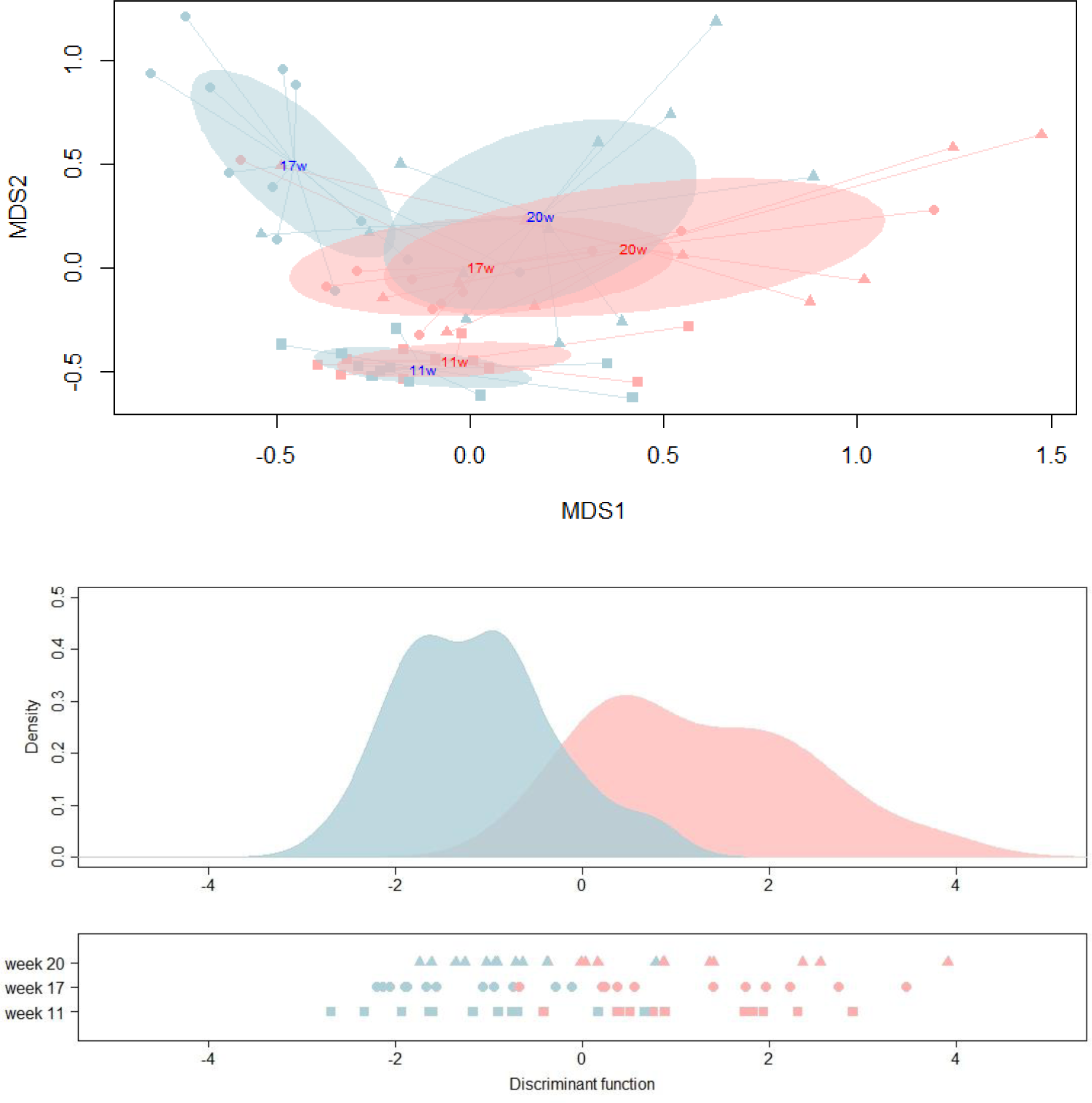
Representation of the Bray-Curtis distance between OTUs using non metric dimensional scaling (stress=0.21), the sampling weeks are indicated in the spidercenter of the corresponding squares, circles and triangles and colored (LRFI line in blue, HRFI in red) (TOP), the discriminant analysis using 98% of the variability via the 23 components of the PCA (MIDDLE) and the position of each animal at each time point on the discriminant axis (BOTTOM).

However, the abundance of *Lactobacillus* OTU25577 and OTU26000 is not enough to predict the pig line from the microbiota composition. Indeed, the relative abundances of nine microbial OTUs were necessary to distinguish LRFI and HRFI pigs with a high specificity (p<0.04, AUC=0.95, Figure S1), while the remaining OTUs marginally improved the discrimination by DAPC applied to the table of abundance of the OTUs of the LRFI and HRFI pig fecal microbiota (p<0.014, AUC=0.9613, Figure 3). Interestingly, each discriminating OTU was found in both LRFI and HRFI pigs, but their individual relative abundance varied from 0.03% to 1.8% (Table S4). In total, the nine discriminating OTUs represented 5% of the total number of sequences in the dataset, further illustrating that the microbial OTUs that differ between the pig lines are not dominant in the gut microbiota.

**Figure 3:**
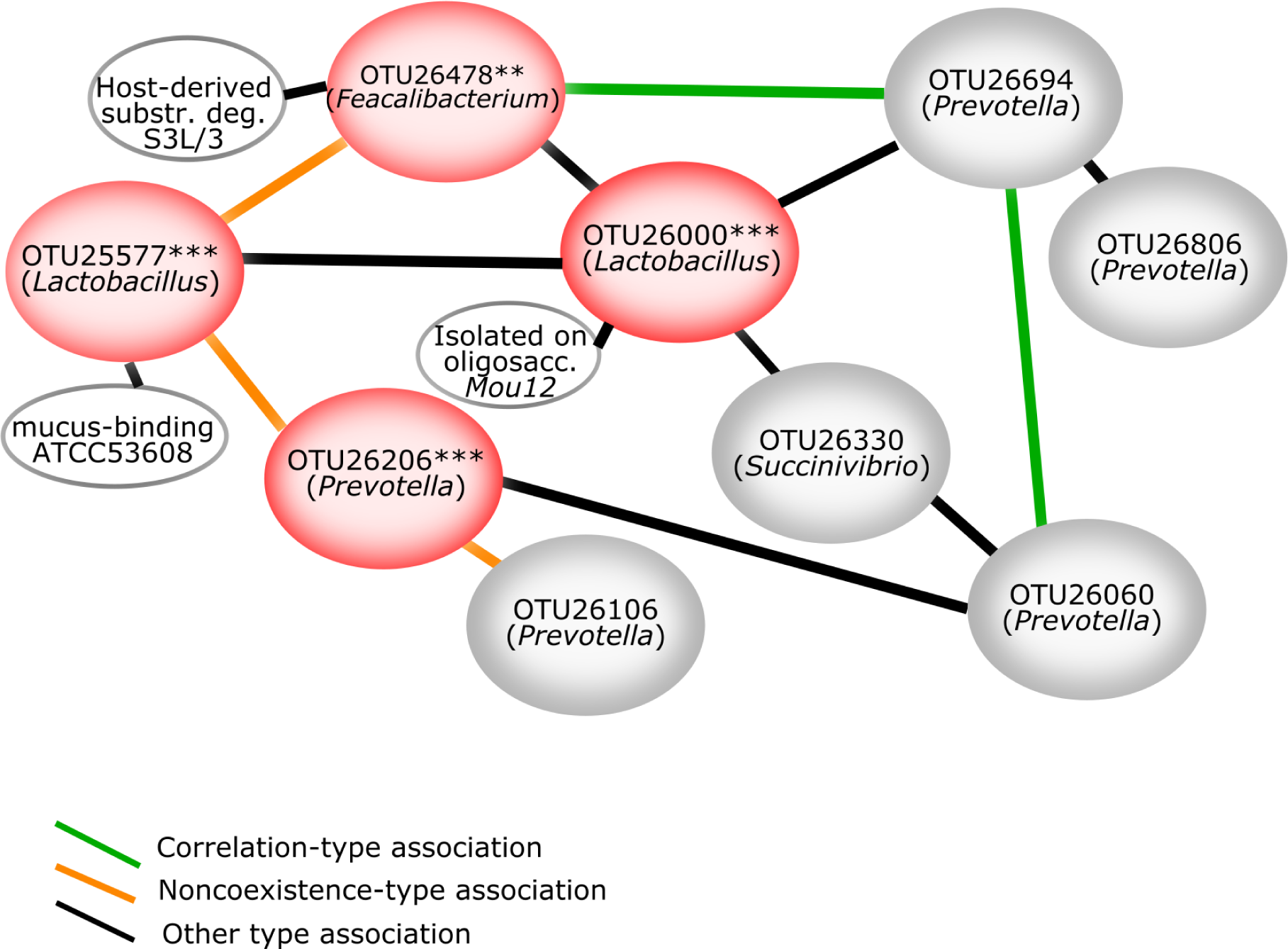
Network of the nine OTUs sufficient to separate the LRFI and HRFI pigs. OTUs in red disks are the OTUs selected via DAPC that are significantly more abundant in high RFI pigs. OTUs in grey disks were relevant to discriminate the lines but not significantly different between lines. When relevant (ID>97%), the closest cultivable relative is reported by the connected white ellipse. Correlations-type relationships are represented with green lines, while orange lines represent exclusion-type relationships. Black lines are relationships that are significant but of neither exclusion- or correlation-types.

Among the nine OTUs that discriminated the LRFI and HRFI lines, four differed significantly according to the non parametric test (Table S4). In addition to the mucus-binding OTU25577, OTU26478 was also depleted in LRFI pigs (0.4 vs 0.6%) and shared 98% similarity with a *Faecalibacterium* from human colon that can utilize host-derived substrates for growth [31]. It is possible that these two OTUs thrive on host-derived substrates, which may be more abundant in HRFI pigs since they have a heavier intestinal tract [10].

Interestingly the nine most discriminant OTUs have complex relationships among each other, including significant noncoexistence relationships that can be detected by the MINE algorithm in gut microbiota [21]. For example, *Lactobacillus*’ OTU25577 has a noncoexistence relationship with *Feacalibacterium*’s OTU26478 (Figure 3), possibly because they share the same niche of host-derived substrates for growth. Other significant nonlinear relationships were detected among the nine most discriminant OTUs (Figure 3) even though their relative abundances were not different between the lines. Our dataset did not show any significant relationship between *Prevotella* and *Ruminococcus* even though their ratio was previously linked to average daily gain [28]. This might be because all animals included in our study would belong to the *Prevotella*-dominated enterotype based on their average percentages of *Prevotella* (44% ± 15%) and *Ruminococcus* (0.3 ± 0.4%). It implies that the variation in ADG in our study is part of the enterotype variation. Some OTUs are improving the discrimination even though they are not significantly different between the groups. This could be a power issue, or it could be a more fundamental flaw in the OTU-approach that cannot cope with functional redundancy. Indeed, an OTU that is replaced by its functional equivalent in half of the samples may not significantly differ between the groups even if the sum of the functional equivalents is relevant. Therefore functional redundancy between OTUs may also represent an inherent limit of the OTU approach.

### Several microbial functions differ between LRFI and HRFI pigs

To overcome the lack of resolution due to the functional redundancy between OTUs we used an alternate approach that aggregates the KEGG orthologs inferred from the sequences of the 16S rRNA gene: a quarter of the 16S rRNA gene sequences (23±9%) mapped to the KEGG reference organisms (corresponding to 43±4% of the OTUs), thereby contributing to an aggregated functional profile. The number of mapped sequences did not differ between the lines (p=0.75). By predicting 6461 KEGG orthologs across the dataset, 898 were significantly different between the lines and linked to KEGG metabolic pathways (Figure S3). Interestingly, 343 metabolic orthologs were overrepresented in LRFI pigs and 555 were overrepresented in HRFI pigs, highlighting pathways that differed between LRFI and HRFI pigs (Figure S2 and S3). For example, the KEGG orthologs (KO) 02517, 03272, 03271, 02535, 02536 and 00748 are all involved in the lipopolysaccharide biosynthesis and overrepresented in LRFI pigs (Figure S2, and suppl table). In contrast, KO 03886, 03887, 03888, 02259, 02301, 02826, 02827 and 02829 suggest that oxidative phosphorylation is higher in HRFI pigs, possibly through the use of the oxygen present in the upper digestive tract [32].

### Aggregated metabolic functions are more predictive than OTUs

Predicting the KOs from the 16S rRNA gene sequence data and aggregating them to get functional community profiles achieved a better separation of the pig lines than the OTU approach (Figure 4, p<0.001, AUC=0.9848). This also seems to be observed in divergent lines of fat and lean chicken using 6739 KOs [33]. Our pig lines could be distinguished using only nine of the 6461 predicted metabolic functions. The DAPC approach highlighted 9 KO that are sufficient to discriminate the lines: they include enzymes (K00656, K01181, K01537), iron transporters (K02014), proteins linked to bacterial motility (K03406) and chemotaxis (K03406), but also transposases (K07484) and symporters (K03310, K03308). Out of the nine functions selected by DAPC, four were significantly different between the lines: the methyl-accepting chemotaxis protein was overrepresented in LRFI pigs while the beta-xylanase, the ATPase and the transposase were underrepresented (Table S5). The better resolution obtained via KOs suggests that the direct transposition of results obtained by the OTU approach to another experiment is challenging. For example, the HMP dataset showed that similar functions could be achieved with very different gut microbial communities [34]. Hence, conversion to KOs or direct shotgun sequencing of functional genes as in the recently published gut microbiome porcine catalogue [35] could provide more generality to the results.

**Figure 4:**
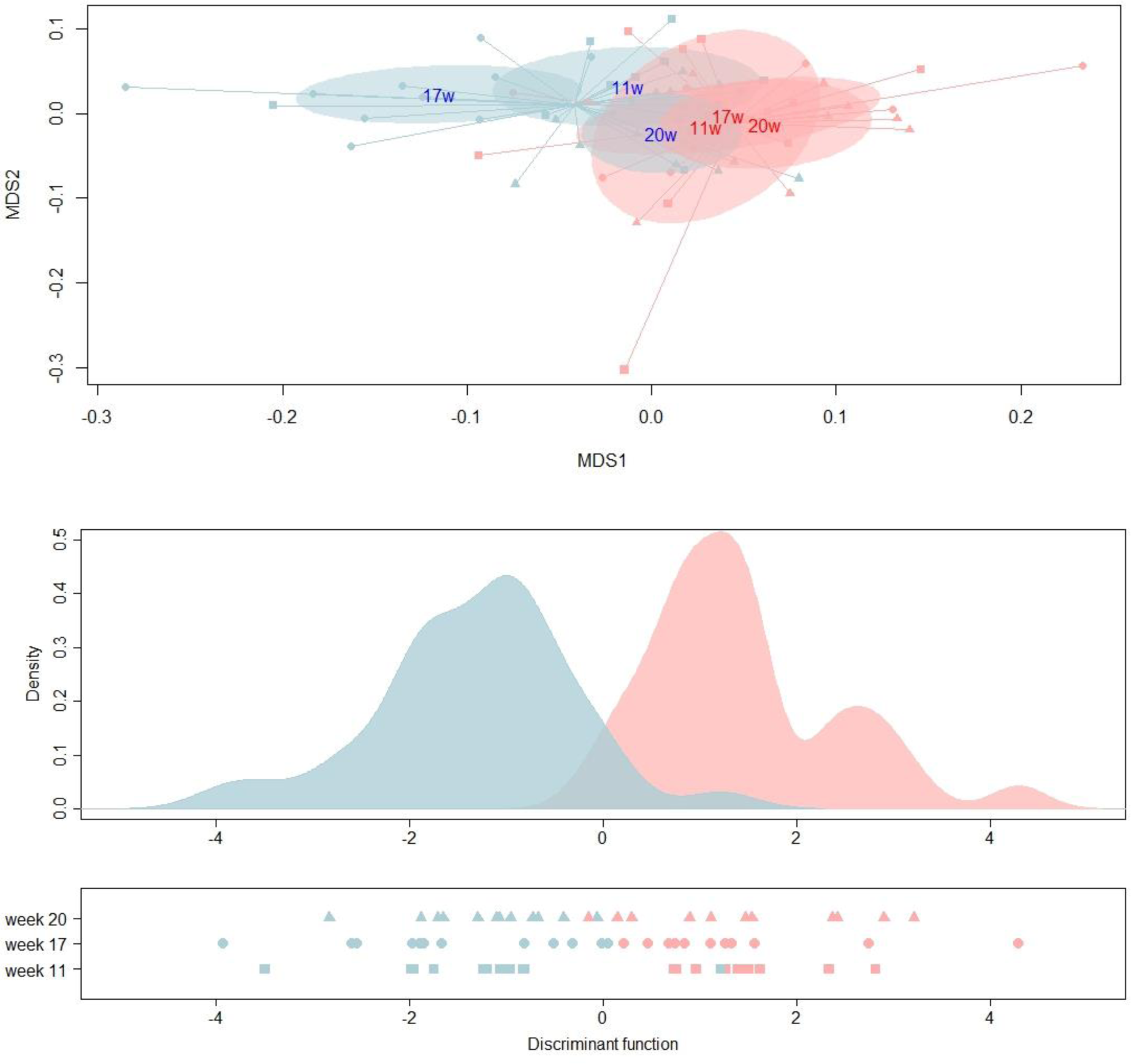
Representation of the Bray-Curtis distance between KOs using non metric dimensional scaling (stress=0.12), the sampling weeks are indicated in the spidercenter of the corresponding squares, circles and triangles and colored (LRFI line in blue, HRFI in red) (TOP); the discriminant analysis using 99% of the variability via PCA (MIDDLE) and the position of each animal at each time point on the discriminant axis (BOTTOM).

### The functional redundancy is higher in LRFI pigs

Interestingly the KO data revealed that LRFI pigs have more metabolic redundancy than their HRFI counterparts (22 ± 17% more OTUs sharing a particular KO in LRFI, p_paired_wilcox =2 10^-16^). Surprisingly almost each metabolic ortholog that was differentially expressed between LRFI and HRFI was shared by at least 2 Silva OTUs (Figure 5). The unique exception was K00608 which is an aspartate carbamoyltransferase that is carried only by an uncultured *Veillonellaceae*. The most shared ortholog was K03639 involved in folate biosynthesis to which 185 and 146 OTUs contributed in LRFI and HRFI pigs, respectively. Functional redundancy of gut microbes is obvious while comparing the ribosomal sequences to the functions observed via shotgun metagenomics [36], yet measuring precisely the functional redundancy and its role in ecosystems functioning remains elusive. Functional redundancy increased the degradation of cellulose in a simplified ecosystem with up to 8 species [37] but the mechanisms at play in our pig lines seem more complicated. Indeed gut microbes might partly explain why the colon of LRFI pigs contained more propionic acid and less branched-chain acids [10], but the lower daily feed intake and the greater ileal digestibility of NDF in LRFI pigs cannot be ruled out. At any rate LRFI pigs have a lower apparent total tract digestibility of NDF [10], which contradicts the hypothesis that the difference in feed efficiency is simply due to microbes giving access to the fibrous part of the feed. In theory, functional redundancy could provide greater guarantees that some will maintain functioning even if others fail, but this theory is more based on converging clues that thorough testing for gut microbial communities [36].

**Figure 5:**
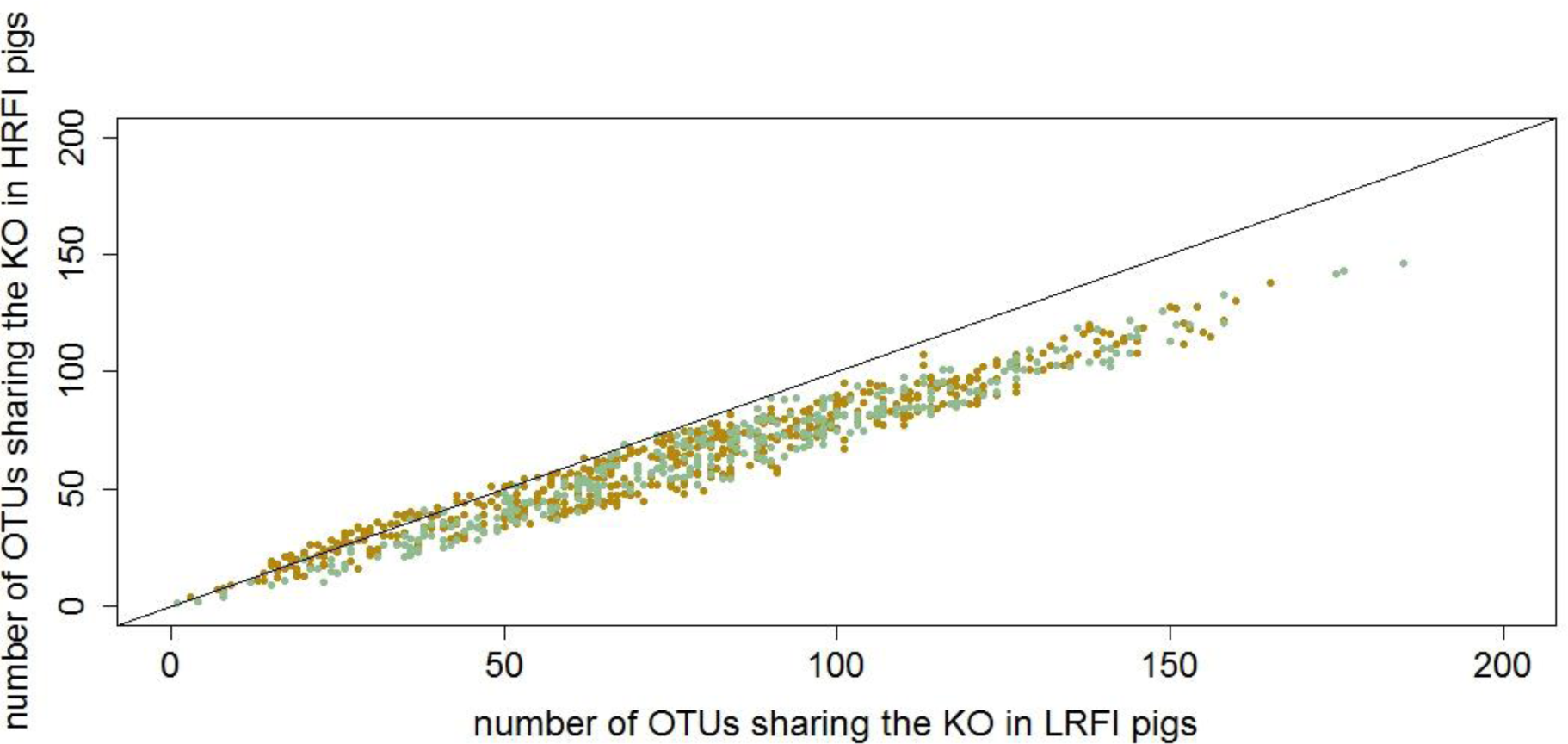
Metabolic redundancy characterized by the number of OTU sharing a particular metabolic KEGG orthologs in LRFI and HRFI pigs. KOs overrepresented in LRFI are in green and KOs underrepresented in LRFI pigs are in yellow.

## Conclusions

In conclusion, LRFI and HRFI pigs have distinct microbial communities in their gut: this is shown by the OTU approach and by the fraction of sequences mapping on the KEGG database for which metabolic functions can be aggregated. Remarkably, almost every function that differ between pig lines are shared between several microbes, which suggests that functional redundancy is the norm rather than the exception even if the Tax4Fun inferences have to be verified by future work. Therefore describing the variation of OTUs seems more difficult to generalize to other studies than describing functional categories. More work is required to determine whether the modified metabolism driven by the host selection resulted in modulated intestinal tract environment and thus the shift of microbiota composition, such as previously observed in mice [38] or whether the gut microbes of LRFI and HRFI pigs trigger the changes of residual feed intake due to their functional redundancy.

## Declarations

### Ethics approval and consent to participate

Not applicable under the French rules for ethics approval because they are part of an on-going selection scheme and no tissue was taken from the animals.

### Consent for publication

Not applicable.

### Availability of data and material

The datasets generated and/or analyzed during the current study are available in the NCBI repository, https://www.ncbi.nlm.nih.gov/bioproject/PRJNA344970.

### Authors’ contributions

HG, CRG and SC designed the study of the microbiota, HG designed the selection experiment, YB performed the selection and sampling of the pigs, CRG and JE extracted the microbial DNA and chose the primers, AC analyzed the microbial sequences, AC and OZ performed the statistical analyses, LC performed the heatmap, OZ and HG drafted the manuscript. All the authors edited the manuscript and approved the paper.

## Acknowledgements

We thank the staff of the GenESI experimental farm for the data and sample collection. We thank Béatrice Gabinaud for the preparation of the libraries. We thank the GeT-PlaGe platform in Toulouse for sequencing the libraries and providing high-quality sequences.

---

**Figure S1:**
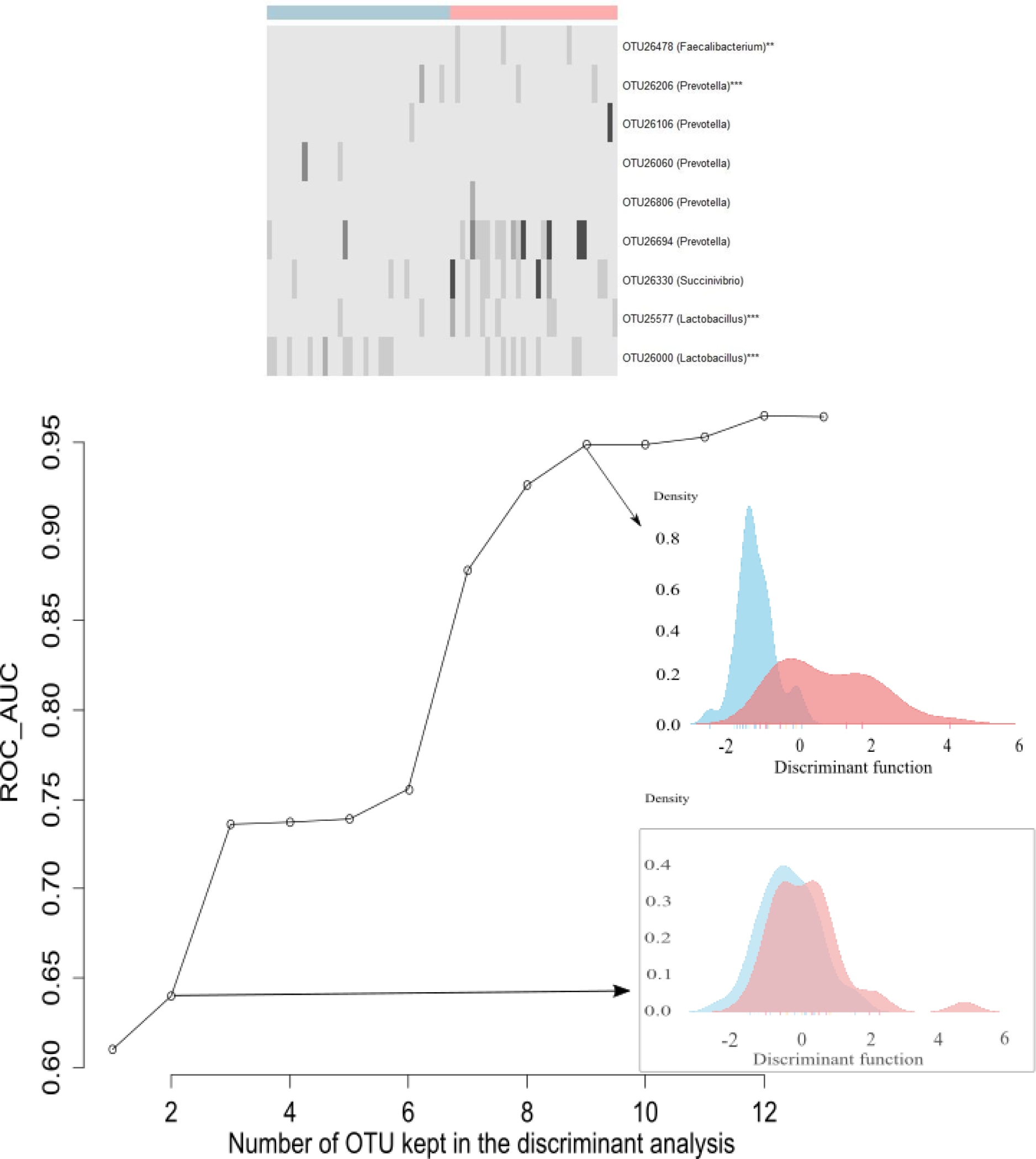
Relative abundance of the discriminative OTUs (TOP) and area under the ROC curve (BOTTOM) to indicate the quality of the separation between the low- and high-RFI lines according of their microbiota OTU composition. The higher inset shows the separation using the top 9 discriminating OTUs (the stars indicate p_kruskal_<0.05). The highest relative abundance of the map is 6% (*Succinivibrio*). The lower inset shows the absence of separation when using 2 OTUs and the improvement when keeping the 9 most discriminative OTUs. In both insets, the density for the low RFI pigs is indicated in blue, whereas the density of the HRFI pigs is indicated in red.

**Figure S2:**
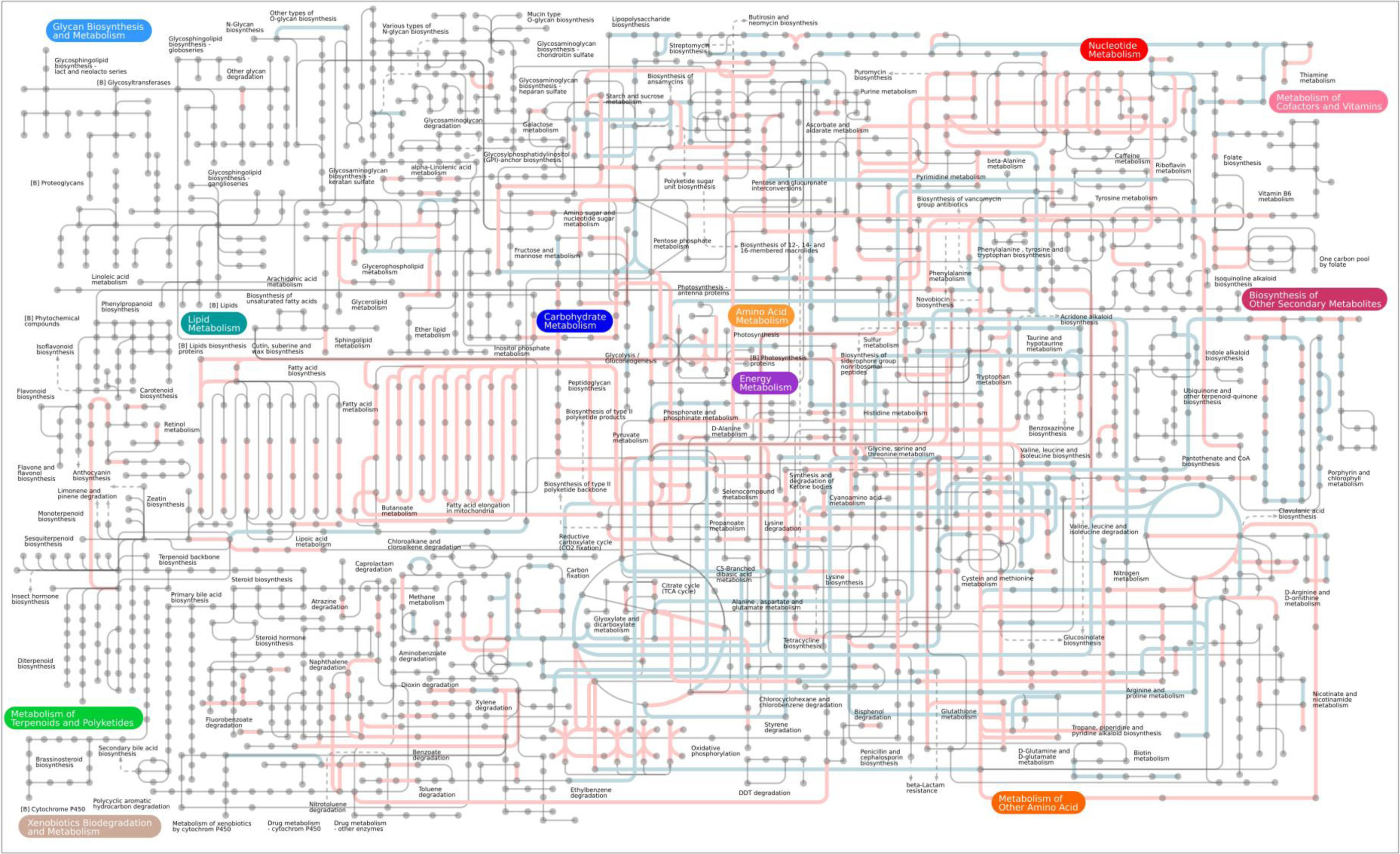
Mapping of the unambiguous KEGG orthologs that are significantly higher in LRFI and HRFI pigs (blue and red, respectively)

**Figure S3:**
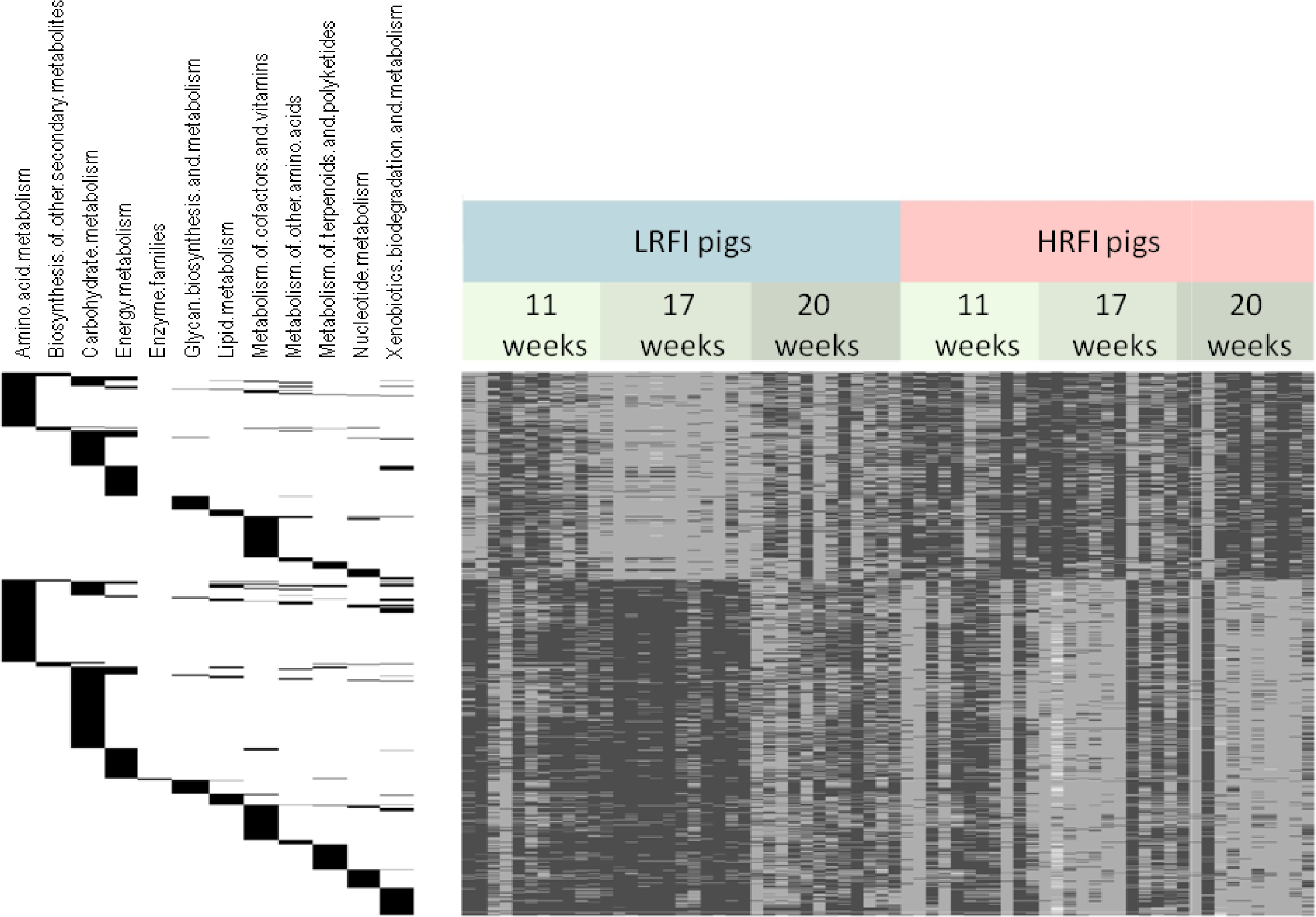
Heatmap of the 898 KEGG orthologs (KO) linked to metabolism that differ between LRFI and HRFI. The left part indicates the appartenance of the KO to a class. The top 343 lines represent the KOs that are overrepresented in LRFI pigs, while the 555 bottom lines represent the KOs that are underpresented in LRFI. The rows are normalized to 1 so that KOs with contrasted abundance can be visualized together.

**Table S1:** composition of the feed.

| **Ingredient** | **g/kg** |
| --- | --- |
| Triticale | 150 |
| Wheat | 240.6 |
| Barley | 100 |
| Wheat bran | 80 |
| Protein peas | 161 |
| Rapeseed meal | 100 |
| Soybean meal | 15 |
| Vegetable oil | 27 |
| Clay | 10 |
| Calcium carbonate | 4.7 |
| Dicalcium phosphate ? | 10 |
| Wheat DDGS | 70 |
| Lysine | 4.7 |
| Methionine | 0.5 |
| Choline | 0.4 |
| Salt | 4 |
| Threonine | 4.3 |
| Mineral and vitamin premix | 10 |
| Belfeed premix powder | 5 |
| Natuphos Premix powder | 2.8 |
| **Analyzed chemical composition** |  |
| DM,% | 88.3 |
| GE, kJ/g DM | 18.61 |

**Table S2:** Average body weights of low RFI (LRFI, N=12) and high RFI (HRFI, N=11) pigs.

| Age | LRFI | HRFI | Significance (p value) |
| --- | --- | --- | --- |
| 11 weeks (kg) | 28.8 ± 3.5 | 29.1 ± 4.5 | 0.87 |
| 17 weeks (kg) | 60.1 ± 6.1 | 65.9 ± 7.2 | 0.05 |
| 20 weeks (kg) | 78.6 ± 7.1 | 85.4 ± 7.2 | 0.03 |

**Table S3:** Average age to reach body weights milestones.

| Weight milestone | Age of LRFI pigs | Age of HRFI pigs | Significance (p value) |
| --- | --- | --- | --- |
| 35 kg | 83.5 days | 81.7 days | 0.52 |
| 65 kg | 122.2 days | 115.7 days | 0.05 |
| 95 kg | 155.2 days | 147.0 days | 0.03 |

**Table S4:** OTUs selected for their discriminative power between LRFI and HRFI pigs.

| #OTU | OTU taxonomy | Mean relative abundance in LRFI | Mean relative abundance in HRFI | P value of the kruskal-wallis test | ID of the representative 16S rRNA gene sequence in the dataset (number of the animal and unique sequence ID) |
| --- | --- | --- | --- | --- | --- |
| 25577 | *Lactobacillus* | 0.03% | 0.44% | 0.0000002 | 11P1134572__HG0G3PF01C1JPK |
| 26000 | *Lactobacillus* | 0.39% | 1.77% | 0.0000018 | 241P1134583__HG0G3PF02GJTRJ |
| 26206 | *Prevotella* | 0.11% | 0.43% | 0.0008 | 113P1135241__HG0G3PF01EGMNB |
| 26478 | *Faecalibacterium* | 0.38% | 0.63% | 0.0049 | 247P1172481__HG0G3PF02JRRG6 |
| 26060 | *Prevotella* | 0.50% | 1.09% | 0.0532 | 246P1135462__HG0G3PF02GUFDK |
| 26694 | *Prevotella* | 0.65% | 0.95% | 0.1170 | 12P1135262__HG0G3PF01DTAG7 |
| 26806 | *Prevotella* | 1.07% | 0.90% | 0.1945 | 237P1135333__HG0G3PF02F1KUE |
| 26330 | *Succinivibrio* | 0.27% | 0.06% | 0.3805 | 117P1162651__HG0G3PF01EASC7 |
| 26106 | *Prevotella* | 0.08% | 0.21% | 0.7676 | 266P1134571__HG0G3PF02FQFSX |

**Table S5:**
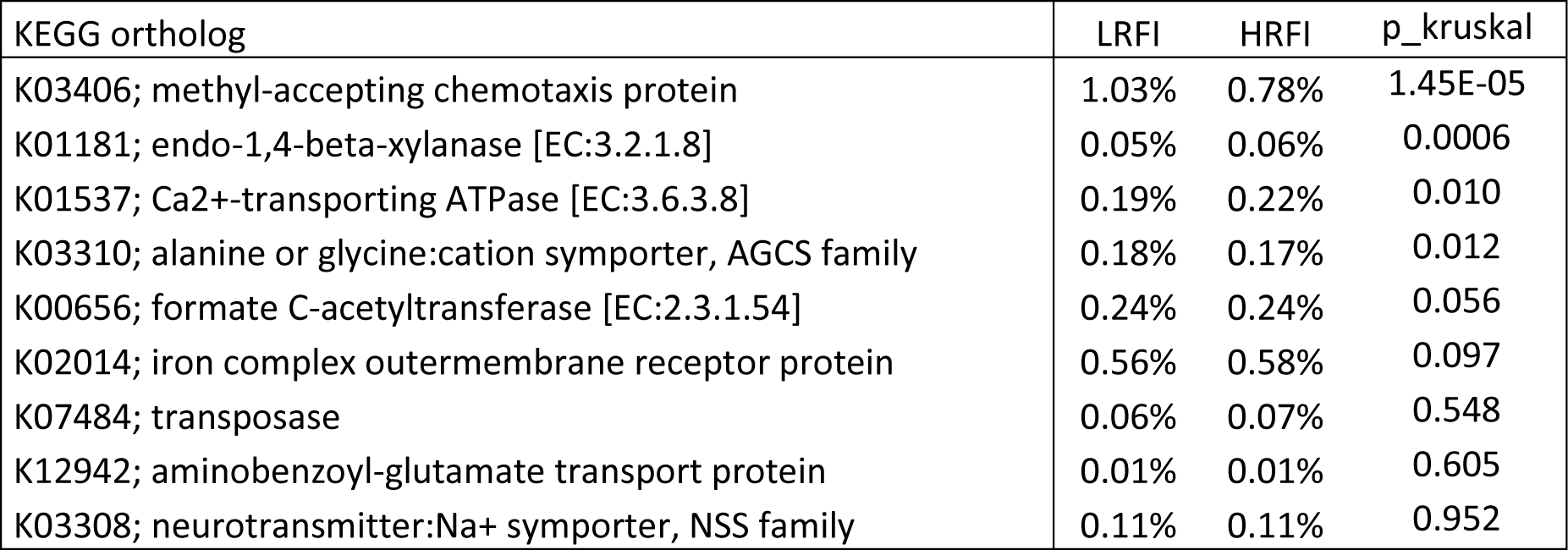
KEGG orthologs selected for their discriminative power between LRFI and HRFI pigs.

| KEGG ortholog | LRFI | HRFI | p_kruskal |
| --- | --- | --- | --- |
| K03406; methyl-accepting chemotaxis protein | 1.03% | 0.78% | 1.45E-05 |
| K01181; endo-1,4-beta-xylanase [EC:3.2.1.8] | 0.05% | 0.06% | 0.0006 |
| K01537; Ca<sup>2+</sup>-transporting ATPase [EC:3.6.3.8] | 0.19% | 0.22% | 0.010 |
| K03310; alanine or glycine:cation symporter, AGCS family | 0.18% | 0.17% | 0.012 |
| K00656; formate C-acetyltransferase [EC:2.3.1.54] | 0.24% | 0.24% | 0.056 |
| K02014; iron complex outer membrane receptor protein | 0.56% | 0.58% | 0.097 |
| K07484; transposase | 0.06% | 0.07% | 0.548 |
| K12942; aminobenzoyl-glutamate transport protein | 0.01% | 0.01% | 0.605 |
| K03308; neurotransmitter:Na<sup>+</sup> symporter, NSS family | 0.11% | 0.11% | 0.952 |

